# ER-embedded UBE2J1/RNF26 ubiquitylation complex in spatiotemporal control of the endolysosomal pathway

**DOI:** 10.1101/2020.10.02.323576

**Authors:** Tom Cremer, Marlieke L.M. Jongsma, Fredrik Trulsson, Alfred C.O. Vertegaal, Jacques J.C. Neefjes, Ilana Berlin

**Author notes:** These authors have contributed equally to this work (Marlieke L.M. Jongsma, Fredrik Trulsson).

## Abstract

The endolysosomal system fulfills a wide variety of cellular functions, many of which are modulated through interactions with other organelles. In particular, the ER exerts spatiotemporal constraints on the organization and motility of endosomes and lysosomes. We have recently described the ER transmembrane E3 ubiquitin ligase RNF26 to control perinuclear positioning and transport dynamics of the endolysosomal vesicular network. We now report that the ubiquitin conjugating enzyme UBE2J1, also anchored in the ER membrane, collaborates with RNF26 in this context, and that the cellular activity of this E2/E3 pair, localized in a perinuclear ER subdomain, is underpinned by transmembrane interactions. Through modification of its substrate SQSTM1/p62, the ER-embedded UBE2J1/RNF26 ubiquitylation complex recruits endosomal adaptors to immobilize their cognate vesicles in the perinuclear region. The resulting spatiotemporal compartmentalization of the endolysosomal system between the perinuclear vesicle cloud and the cell periphery facilitates timely downregulation of endocytosed cargoes, such as EGFR.

## Introduction

Eukaryotic cells have evolved a complex architecture encompassing the nucleus, cytoplasm and various specialized organelles, all confined within a small three-dimensional space. While compartmentalization enables cells to maintain order, interactions between compartments in turn offer opportunities for integration and coregulation of essential cellular processes. For instance, the ER, typically the cell’s largest organelle, offers an excellent platform for supervision of smaller intracellular structures. In fact, membrane contact sites between the ER and virtually every intracellular organelle have been reported to date, allowing controlled exchanges of information and materials to occur between them (Phillips & Voeltz, 2016, Wong, Gatta et al., 2019). Uncovering ways in which the ER communicates with and influences other organelles is crucial to our understanding of how cells coordinate their internal affairs and respond to their environment.

The endolysosomal system, comprised of a vesicular network whose members are both physically independent and functionally interconnected, presents a unique case in its relationship with the ER (Raiborg, Wenzel et al., 2015b). The endocytic compartment fulfills a wide variety of cellular roles, ranging from regulation of signaling and proteostasis (Di Fiore & von Zastrow, 2014, Khaminets, Behl et al., 2016) to control of cell polarity (Jewett & Prekeris, 2018), migration (Malinova & Huveneers, 2018, Paul, Jacquemet et al., 2015), defense against pathogenic invaders (Taguchi & Mukai, 2019) and communication between cells (Maas, Breakefield et al., 2017). Once nascent endosomes bud inwards from the plasma membrane, engulfing extracellular milieu, they embark on a journey of maturation, guided in part by the ER (Bakker, Spits et al., 2017). Travelling deeper into the cell interior, endosomes progressively acquire late characteristics of acidity and proteolytic potential (Huotari & Helenius, 2011) and engage in more frequent and persistent contacts with the ER membrane (Friedman JR, 2013). These interactions have been shown to influence endosome localization and motility (Jongsma, Berlin et al., 2016, Raiborg, Wenzel et al., 2015a, Rocha, Kuijl et al., 2009b) and control core processes pertaining to endosome physiology, including cargo sorting (Dong, Saheki et al., 2016, Eden, Sanchez-Heras et al., 2016, Eden, White et al., 2010) as well as membrane tethering, fusion and fission events (Hoyer, Chitwood et al., 2018, Levin-Konigsberg, Montano-Rendon et al., 2019, Rowland, Chitwood et al., 2014, van der Kant, Fish et al., 2013, Wijdeven, Janssen et al., 2016). The rapidly growing diversity in ER-endosome contacts underscores both the importance and complexity of the dialogue occurring between these organelles.

It is becoming increasingly clear that specific functional states of endocytic organelles are connected to their intracellular location (Jia & Bonifacino, 2019, Johnson, Ostrowski et al., 2016, Korolchuk, Saiki et al., 2011)—an attribute strongly influenced by ER-endosome interactions (Neefjes, Jongsma et al., 2017). Under normal circumstances, the bulk of endosomes and lysosomes congregates in a perinuclear cloud around the microtubule-organizing center (MTOC). While many endosomes and lysosomes participating in this cloud tend to exhibit limited motility (Jongsma et al., 2016), some become subject to fast transport to and from the cell periphery (Bonifacino & Neefjes, 2017). This bilateral organization between the perinuclear and peripheral regions of the cell appears critical for efficient maturation of endosomes and, consequently, timely degradation of cargos, such as EGFR (Jongsma et al., 2016). Perinuclear retention of early, late, and recycling endosomes, as well as lysosomes and vesicles of the trans-Golgi network (TGN) is governed by the ER-located ubiquitin ligase RNF26—an integral multimembrane-spanning protein featuring a cytoplasmically exposed RING domain (Jongsma et al., 2016, Qin, Zhou et al., 2014). RNF26 is concentrated predominantly in the perinuclear segment of the ER membrane, which corresponds with its ability to position all endosomal vesicles near the nucleus (Jongsma et al., 2016). Catalytically competent RNF26 attracts and ubiquitylates SQSTM1/p62, a cytosolic ubiquitin adaptor also implicated in selective autophagy (Lamark, Svenning et al., 2017), and the resulting ubiquitin-rich complex then recruits various endosomal adaptors capable of ubiquitin recognition to dock at the ER (Jongsma et al., 2016). How RNF26 activity is regulated to fulfill this role is unknown.

Ubiquitylation, orchestrated by a hierarchical enzymatic cascade (Pickart, 2001), is pervasive in endosome biology (McCann, Scott et al., 2016, Polo, 2012, Raiborg & Stenmark, 2009). In order to become biologically useful, ubiquitin must first be activated by an E1 enzyme. Next, an E2 enzyme receives this activated ubiquitin and can either pass it on to an independent E3 enzyme (as in the case of the HECT family of E3 ligases) or join forces with a RING-type E3 to directly mediate transfer of ubiquitin to a substrate of choice (Stewart, Ritterhoff et al., 2016). While in mammals only 2 E1 enzymes for ubiquitin are known, roughly 40 E2 conjugating enzymes and over 600 E3 ligases have been identified (Zheng & Shabek, 2017). This implies that E2 enzymes usually support multiple E3 ligases, and a given E2 is likely to be involved in diverse biological processes (Gundogdu & Walden, 2019). Ultimately, the type and extent of modification produced by a given E2/E3 pair determines the substrate’s resulting functional state (Kwon & Ciechanover, 2017).

A key missing piece in understanding ubiquitin-regulated positioning of vesicles by the RNF26-associated system is the contribution of a cognate E2 enzyme. Here we identify UBE2J1 as the conjugating enzyme collaborating with RNF26 in the regulation of the perinuclear endosome cloud. We find that an intramembrane interaction between RNF26 and UBE2J1 is necessary for successful assembly of this enzyme complex within a perinuclear ER subdomain. Through ubiquitylation of SQSTM1, and consequent vesicle adaptor recruitment onto the positioning complex, UBE2J1 controls the integrity of the endolysosomal cloud. Hence, UBE2J1 function, like that of RNF26, promotes ligand-mediated trafficking of activated receptors towards acidified compartments, ensuring their timely down-regulation. These findings uncover a new role for UBE2J1, an E2 extensively implicated in ER-associated protein degradation (ERAD) (Burr, Cano et al., 2011, Hagiwara, Ling et al., 2016, Lenk, Yu et al., 2002), and in this light open doors to possible interplay between ERAD and the perinuclear endolysosomal cloud.

## Results

### Depletion of UBE2J1 scatters the perinuclear cloud

Given that the E3 ligase RNF26 employs ubiquitylation to position endosomes and lysosomes, there must also be a collaborating E2 enzyme. To identify E2 ubiquitin conjugating enzyme(s) participating in the formation and maintenance of the perinuclear endosomal cloud, we performed an siRNA-based screen for all known E2 enzymes in the human melanoma MelJuSo cell line. siRNAs inducing dispersion of late endosomes throughout the cytoplasm, and/or accumulation of these organelles at the tips of cells, were selected. Silencing of ubiquitin E2 enzymes UBE2G2, UBE2I, UBE2L6, UBE2N, UBE2J1, UBE2R1, UBE2V1, and UBE2Z, as well as ubiquitin-like conjugating enzymes UFC1 and ATG3 (Stewart et al., 2016), perturbed perinuclear accumulation of endolysosomes marked by the major histocompatibility class II (MHC-II) receptor (Fig. S1A). Among the aforementioned E2 hits, silencing of UBE2J1 afforded the most striking relocalization of the MHC-II^+^ compartment (Fig. S1A), and we therefore focused subsequent validation on UBE2J1 as a candidate E2 for RNF26.

We have previously shown that the perinuclear cloud harbors the entire endosomal pathway, including TGN-derived vesicles, early and late endosomes and lysosomes, and that loss of RNF26 disturbs their localization (Jongsma et al., 2016). We therefore examined whether this phenotype extends to other endosomal structures. Depletion of UBE2J1 phenocopied that of RNF26, resulting in scattering of early (EEA1^+^), recycling (TfR^+^) and late endosomes (CD63^+^), as well as lysosomes (LAMP1^+^) and vesicles of the TGN (M6PR^+^) throughout the cytoplasm (Figs. 1A, B and S1B-E). By contrast, depletion of its closest homologue UBE2J2 did not appear to disrupt endosomal organization (Figs. 1A, B and S1F).

**Fig. 1:**
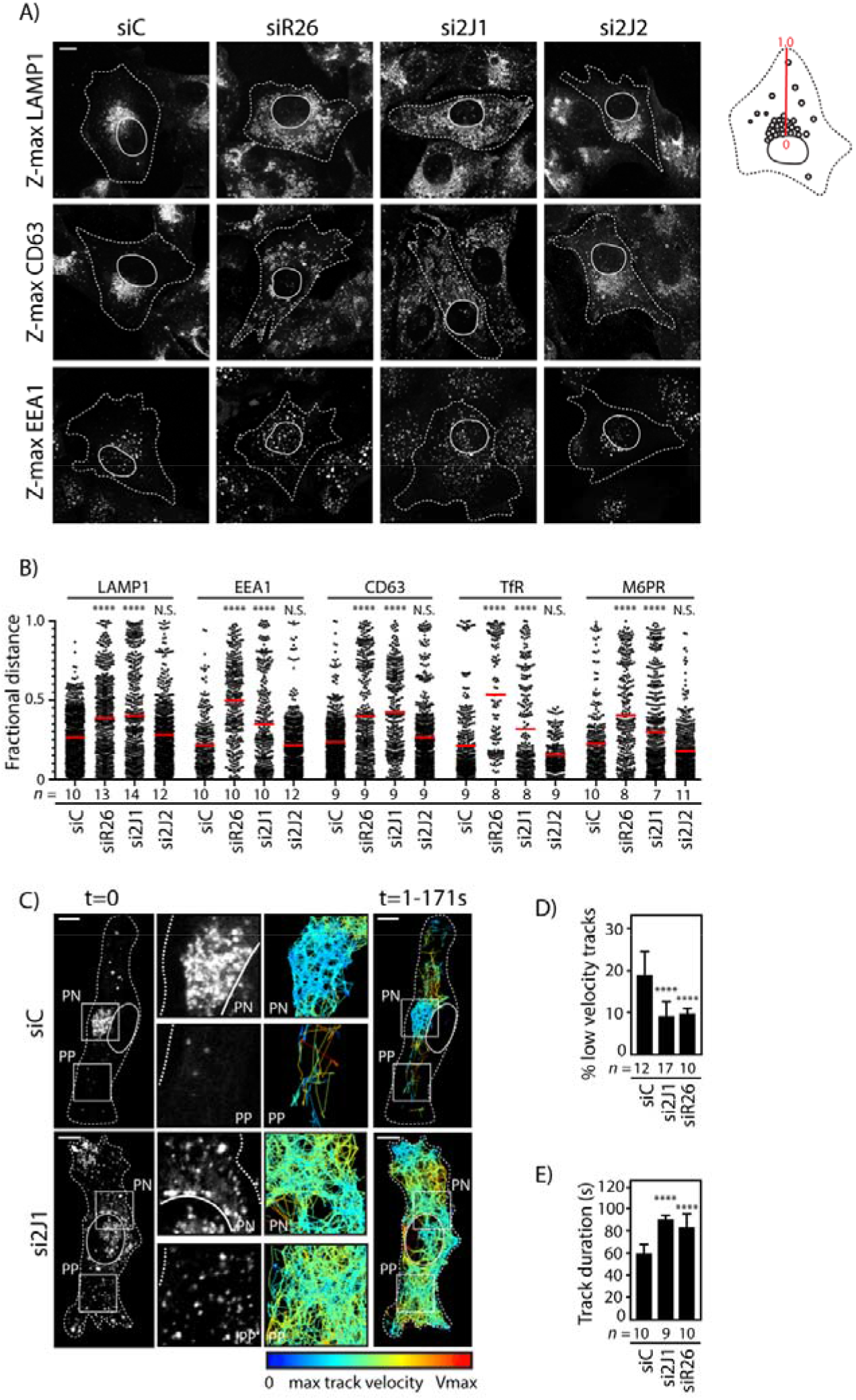
UBE2J1 is required for integrity of the endolysosomal cloud. **A)** Distribution of endosomes and lysosomes in response to depletion of UBE2J1. Representative confocal z-projections of fixed MelJuSo cells transfected with siRNAs targeting UBE2J1 (siRNA# 1), UBE2J2 or RNF26 and immunostained against markers for lysosomes (LAMP1), early endosomes (EEA1), or late endosomes (CD63) are shown. **B)** Vesicle localization expressed as fractional distance of fluorescent pixels along a straight line drawn through the endosomal cloud from the nuclear edge (0) to the plasma membrane (1.0) analyzed from samples in (1A) and (S1B). UBE2J2 depletion inversely affected TfR and M6PR distribution (** and ***, respectively). Red line: mean, n=2 independent experiments. **C)** Organization and dynamics of late compartments as a function of UBE2J1 depletion. Left panels: representative confocal images of live MelJuSo cells transfected with either control siRNA oligo (siControl) or oligo #1 targeting UBE2J1 and treated with LysoTracker FarRed taken at the start of time-lapse, t=0. Original movie is supplemented online. Right panels: vesicle displacement rates depicted on a rainbow color-scale (blue: immobile; red: maximum mobility per time interval) tracked over 171 s at 1.62s per frame using the TrackMate plugin for Fiji. Cell and nuclear boundaries are depicted using dashed or continuous lines, respectively; boxed zoom-ins highlight select perinuclear (PN) and peripheral (PP) regions. **D-E)** Quantification of vesicle motility in control MelJuSo cells (siC) or those depleted of either UBE2J1 (si2J1-1) or RNF26 (siR26). Bar graph reports mean percentage of low velocity tracks (as defined in S1G) per cell (D) or total track duration (E). n=2 independent experiments. Shown is mean+SD. Cell and nuclear boundaries are demarcated using dashed and continuous lines, respectively. Magnification identical for all images. Number of cells analyzed per condition is given under each bar/scatter. Scale bars = 10μm. All statistical significance tested with Students’ T-test. ** = p<0,01; *** = p<0,001; **** = p<0.0001.

The architecture of the endolysosomal system is intimately connected to the motility of its individual vesicles, and dissociation of the perinuclear cloud leads to disordered vesicle transport throughout the cell (Jongsma et al., 2016, Sapmaz, Berlin et al., 2019). We therefore tested whether depletion of UBE2J1 would affect not only the position but also movement of endolysosomes. Under control conditions, acidified compartments (marked by Lysotracker) displayed bimodal motility, with the majority of perinuclear (PN) endolysosomes exhibiting restricted movement relative to a smaller pool of their far more dynamic peripheral (PP) counterparts (Fig. 1C, movie 1). Silencing UBE2J1 abrogated this spatiotemporal distinction, releasing vesicles normally retained in the PN cloud for transport (Fig. 1C, movie 1). As a result, an overall increase in vesicle movement was observed, resembling the condition of RNF26 deficiency (Figs. 1D, E and S1G, H). The E2 enzyme UBE2J1 thus recapitulates the phenotype of RNF26 in imposing spatial and temporal constraints on the endolysosomal system.

### Transmembrane interactions underpin UBE2J1/RNF26 complex in the juxtanuclear ER subdomain

In order for UBE2J1 to act as an E2 enzyme for RNF26, the two proteins must form a complex at the ER membrane. Therefore, we examined whether RNF26 and UBE2J1 colocalize and interact. RNF26 is a multipass transmembrane protein (Qin et al., 2014) and locates in the perinuclear subdomain of the ER in order to restrict the endolysosomal system at the corresponding location in the cytosol (Jongsma et al., 2016). On the other hand, UBE2J1 harbors a single transmembrane domain and distributes all along the ER, as indicated by a high degree of colocalization with the ER protein VAP-A (Fig. 2A, B) (Yang M., 1997). Strikingly, co-expression of RNF26 focused UBE2J1 into the perinuclear ER, resulting in colocalization between the two enzymes (Fig. 2B-D). A similar effect was observed in the presence of a catalytically inactive RNF26 point mutant I382R, but not with RNF26 lacking the RING domain (Fig. 2B-D). Hence, the RING domain of RNF26, critical for its perinuclear localization (Jongsma et al., 2016), also influences the location of UBE2J1.

**Fig. 2:**
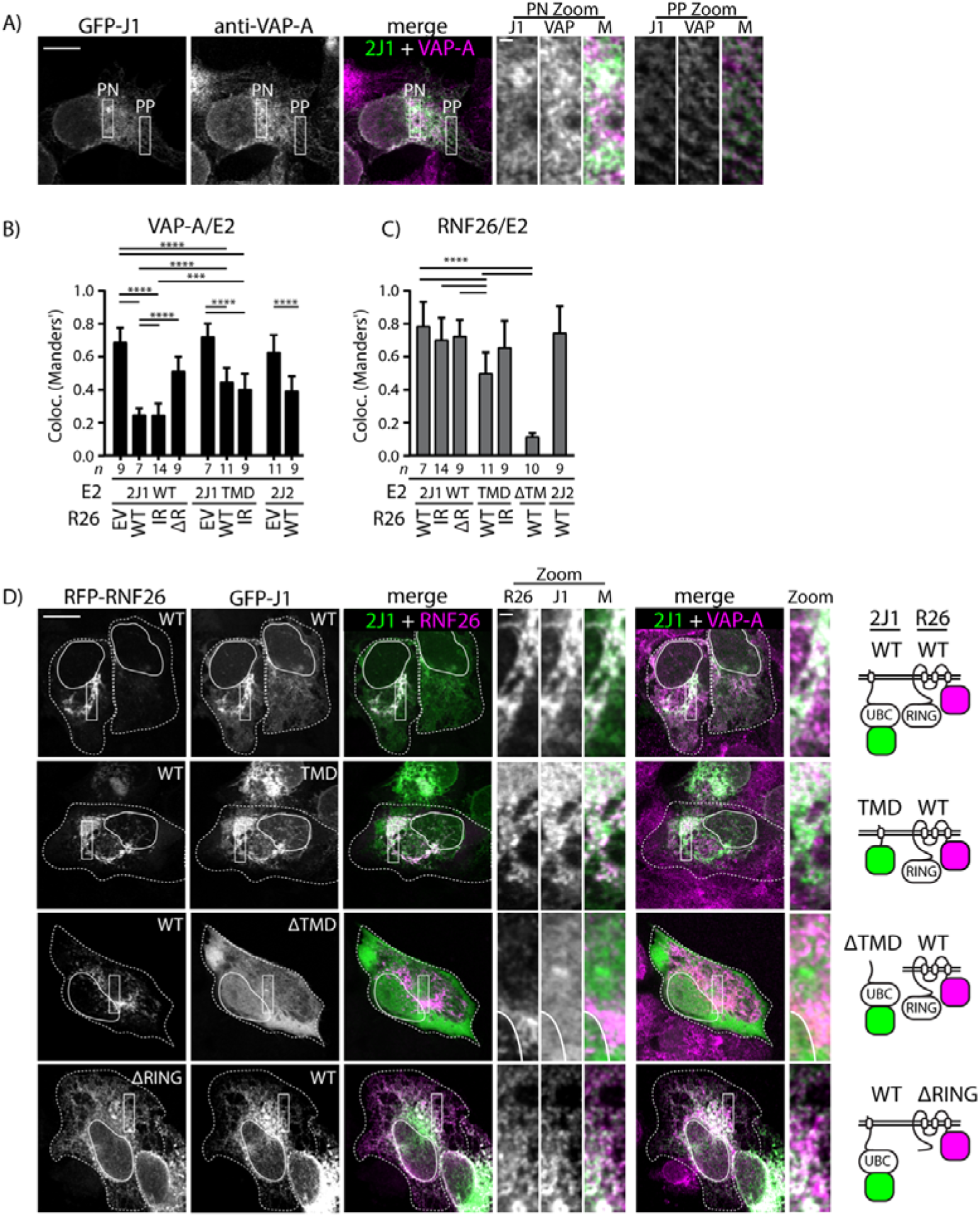
RNF26 recruits UBE2J1 to a perinuclear ER subdomain. **A)** Localization of UBE2J1 in the ER. Representative confocal image of MelJuSo cells expressing GFP-UBE2J1, immunostained against VAP-A (ER marker). Rectangular zoomins highlight select perinuclear (PN) and peripheral (PP) regions. **B)** Colocalization (Manders’ overlap) between either GFP-UBE2J1 WT, -TMD or -UBE2J2 and ER marker VAP-A in single transfected cells (EV) or in the presence of RFP-RNF26 WT, inactive mutant I382R or ΔRING truncation mutant in MelJuSo cells, as indicated. Shown is mean + SD. **C)** Colocalization (Manders’ overlap) between either GFP-UBE2J1 WT, -TMD, -ΔTMD, -UBE2J2 and RFP-RNF26 WT, -IR or -ΔRING truncation mutant in MelJuSo cells, as indicated. Shown is mean + SD. **D)** Localization of UBE2J1 as a function of RNF26. Representative single plane confocal images of MelJuSo cells co-expressing full length GFP-UBE2J1, its transmembrane domain (TMD), or a mutant lacking the TMD (ΔTMD) with either full length RFP-RNF26 or its mutant lacking the RING domain (ΔRING). Immunostaining against VAP-A was used as an ER marker. Overlays of either UBE2J1 and VAP-A or RNF26 are shown and schematic representations of the used constructs are depicted on the right. Rectangular zoom-ins highlight select regions of indicated channels. Scale bar = 10μm. Zoom scale bar = 1μm. Magnification identical for all images. ImageJ mean filter (1 pxl) was used to smoothen images. Cell and nuclear boundaries are demarcated using dashed and continuous lines, respectively. Number of cells analyzed per condition is given under each sample group. All statistical significance tested with Students’ T-test. *** p<0.001; **** p<0.0001.

We further examined the requirements for complex formation between UBE2J1 and RNF26. Endogenous UBE2J1 readily co-precipitated with ectopically expressed wild type RNF26, as well as its mutants I382R and ΔRING (Fig. 3A), indicating that the RING domain, while important for specifying intracellular location of this enzyme complex, is not necessary for its formation. These observations suggested that UBE2J1 and RNF26 may interact via their respective trans-membrane domains (TMDs) instead. Indeed, the single TMD of UBE2J1, which on its own took residence throughout the ER membrane (Fig. S2A), could be drawn into the perinuclear region by RNF26 (Fig. 2B-D), although to a lesser degree than the full length UBE2J1 (Fig. 2B, C). In contrast, UBE2J1 lacking its TMD did not localize to RNF26, but remained dispersed throughout the cytosol (Fig. 2B-D). In line with these observations, RNF26-ΔRING co-isolated with the TMD of UBE2J1, but not its TMD-deficient soluble fragment (Fig. 3B). This suggests that UBE2J1 binds RNF26 in a perinuclear ER subdomain, and that this interaction relies on the respective transmembrane determinants of these two proteins. In addition to UBE2J1, RNF26 was also found to interact and colocalize with UBE2J2 (S2B, C), loss of which did not influence intracellular localization of the endolysosomal system (Fig. 1A, B). We therefore tested whether UBE2J1—but perhaps not UBE2J2—shares RNF26’s substrate SQSTM1, as described below.

**Fig. 3:**
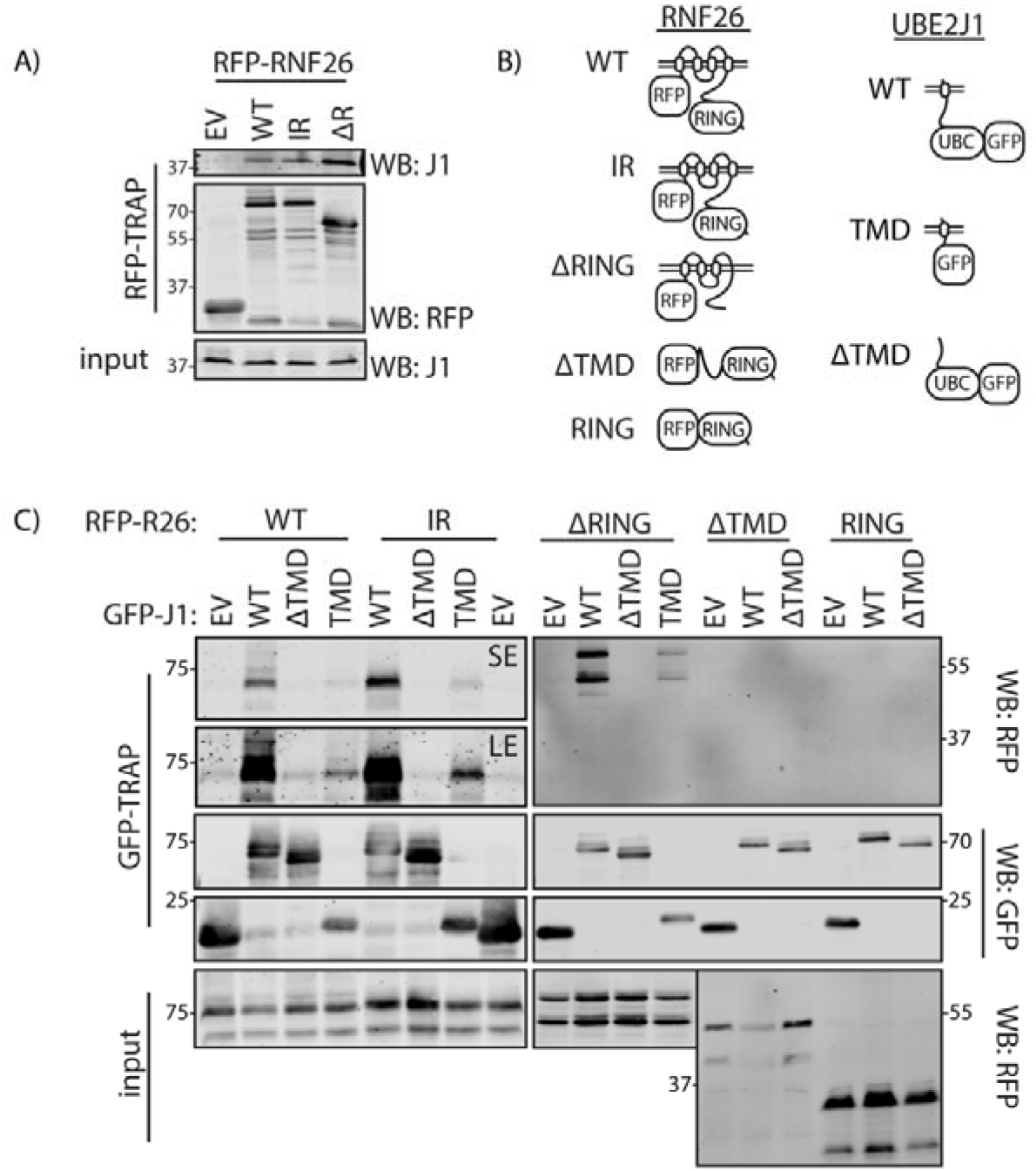
Interaction between RNF26 and UBE2J1 requires their TMDs. **A)** Interaction of overexpressed RFP-RNF26, IR mutant, or ΔRING mutant with endogenous UBE2J1 as assayed by CoIP. Cell lysates were used as input control. Position marker proteins indicated. **B)** Schematic representation of the UBE2J1 and RNF26 constructs used in these interaction studies. **C)** Interactions between RNF26, its inactive mutant -I382R (IR), -ΔRING, -RING, or - ΔTMD truncations mutant versus UBE2J1 WT, -TMD, or -ΔTMD as indicated, assayed by co-IP in HEK293 cells. For WT and IR sample groups, four times as much sample was loaded and enhanced signal images are also shown (LE). For input samples in RNF26 WT, - IR and -ΔRING sample groups, post-IP lysate was subjected to a secondary IP with an excess of RFP-TRAP beads, while post-nuclear lysates were used to depict input levels in RNF26 RING or ΔTMD sample groups. A representative result from duplicate experiments. Position marker proteins indicated.

### UBE2J1 mediates ubiquitylation of SQSTM1 for recruitment of vesicle adaptors to RNF26

While UBE2J1 interacts with RNF26, this does not imply that the catalytic activity of the E2 enzyme is involved in RNF26-mediated endosome positioning. To this end, we created UBE2J1 knockout HeLa cells using CRISPR/Cas9 gene editing and aimed to reverse the UBE2J1-depleted endosomal phenotype by reintroducing the enzyme herein. Similar to siRNA mediated depletion (Fig. 1A), UBE2J1 knockout cells featured a dispersed CD63^+^ late endosomal compartment (Fig. 4A, B). Re-expression of wild type UBE2J1 in this setting, but not its catalytically dead mutant UBE2J1-C91S, facilitated centering of late endosomes in the perinuclear area (Fig. 4C, D). This implies that a functional UBE2J1 enzyme is required for perinuclear localization of the late endosomal compartment. We then wondered whether a functional RNF26/UBE2J1 complex would accumulate ubiquitinate species inside the cell. While the combination of RNF26 and catalytically competent UBE2J1 stimulated deposition of ubiquitylated species onto the E2/E3 complex, co-expression of either inactive UBE2J1 or wild type UBE2J2 nearly abolished ubiquitylation at RNF26-positive sites (Fig. 4E, F). Next, we tested whether UBE2J1 can mediate ubiquitin modification of RNF26 substrate, SQSTM1. Ubiquitylation of SQSTM1 was markedly enhanced by overexpression of catalytically competent UBE2J1, similar to the effect of RNF26 (Fig. 4G). This was not observed in response to overexpression of UBE2J2 (Fig. 4G), indicating that even though UBE2J2 can interact with RNF26, the two do not share SQSTM1 as a substrate. However, fusing the catalytic domain of UBE2J1 to the TMD of UBE2J2 (2J1(J2TMD)), still produced an ER-located enzyme competent of modifying SQSTM1 (Figs. 4G and S3). On the contrary, a similar chimera harboring the TMD of MOSPD2 (2J1(MSD2TMD)), an unrelated single-spanning protein anchored in the ER membrane, did not afford ubiquitylation of SQSTM1 (Figs. 4G and S3). These results imply that appropriate catalytic and TMD determinants are required on the part of the E2 to make a productive pair with RNF26 and suggest that a high degree of E2 selectivity operates in the pathway(s) responsible for ubiquitylation of SQSTM1.

**Fig. 4:**
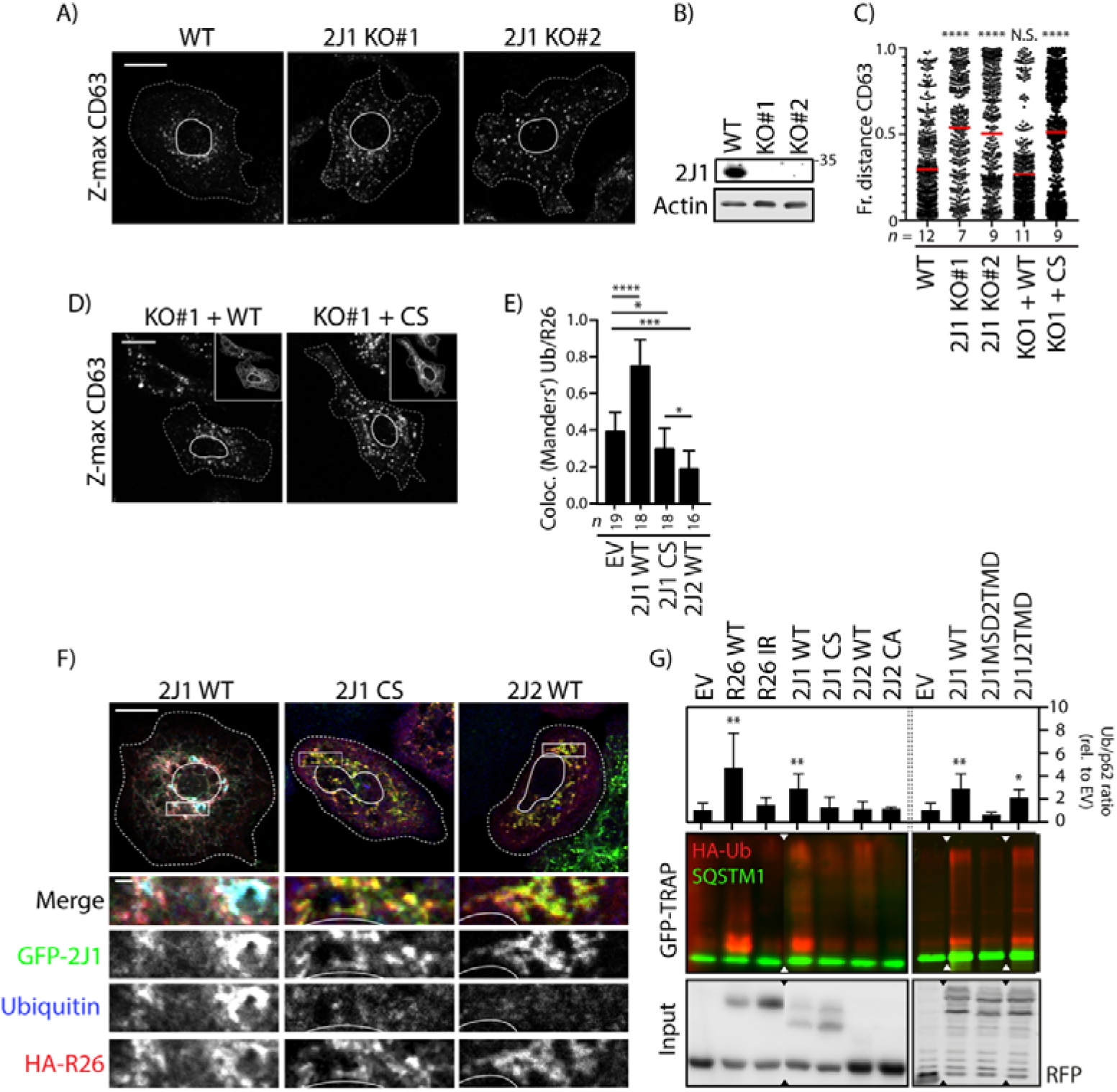
UBE2J1 activity is required for RNF26 function. **A)** Late endosome distribution in WT HeLa cells or two UBE2J1 KO clones. Representative maximum projection confocal images of CD63 immunostaining is shown. **B)** WB validation of UBE2J1 KO in two clonal cell lines created by CRISPR-Cas9 vector transfection and limiting dilution. Actin was used as loading control. Position protein marker indicated. **C)** Vesicle localization depicted as fractional distance as in Figure 1C of samples in (A) and (C). Results from two independent experiments, significance was tested versus WT sample. **D)** Late endosome distribution in HeLa UBE2J1 KO cells expressing either GFP-UBE2J1 WT or inactive -CS. Maximum confocal Z-projections of cells immunostained against CD63. **E)** Colocalization (Manders’ overlap) between Ubiquitin and HA-RNF26, sampled from (E), n=3 independent experiments. **F)** Ubiquitin recruitment to RNF26/UBE2J1-positive structures as a function of UBE2J1 activity. Representative confocal images of HeLa cells expressing HA-RNF26 and either GFP control, GFP-UBE2J1 WT or -CS (green). Cells were immunostained for HA (red) and Ubiquitin (blue). Representative single focal plane fluorescence overlays and single channel zooms are shown in black-white and a colored merge panel. **G)** Ubiquitylation of SQSTM1 as a function of RNF26, UBE2J1 or UBE2J2 activity. GFP-SQSTM1 (green) was immunoprecipitated from cells co-expressing HA-Ubiquitin (red) and either RFP-RNF26 versus mutant IR, RFP-UBE2J1 versus mutants CS, MSD2TMD, 2J2TMD or RFP-UBE2J2 versus mutant CS or vector control (EV). Input shows total cell lysate. Triangles indicate where lanes were excised from the original scan. GFP-SQSTM1 signal was adjusted for comparison. Position of molecular weight standards indicated. Relative amounts of HA-Ub conjugated to GFP-SQSTM1 in each condition were quantified and normalized to EV. Scale bars = 10μm. Zoom scale bar = 1μm. Magnification identical for all images. Cell and nuclear boundaries are demarcated using dashed and continuous lines, respectively. Number of cells analyzed per condition is given under each sample group. All statistical significance tested with Students’ T-test. * p<0.05; ** p<0.01; *** p<0.001; **** p<0.0001. Shown is mean + SD from triplicate experiments.

Next, having identified UBE2J1 as a compatible partner for RNF26, we set out to examine whether this E2 enzyme plays an active role in the recruitment of vesicle adaptor proteins to the RNF26-positioning complex. As a consequence of RNF26 enzymatic action, SQSTM1 is ubiquitinated for recognition by a number of endosomal membrane adaptors that contain a ubiquitin-binding domain. These adaptors include EPS15 and TOLLIP, which link early and late endosomes, respectively, to the ER-embedded positioning complex (Jongsma et al., 2016). Perturbation in the cognate E2 activity should then dissociate such ubiquit-independent bridges between the ER and endosomes. As expected, ectopic expression of inactive, but not wild-type, UBE2J1 strongly diminished these contacts, as judged by fluorescence signal overlap of either TOLLIP or EPS15 (Fig. 5A-C) with RNF26. These observations further support a pivotal role of the ubiquitin conjugating activity of UBE2J1 in positioning of the endosomal pathway.

**Fig. 5:**
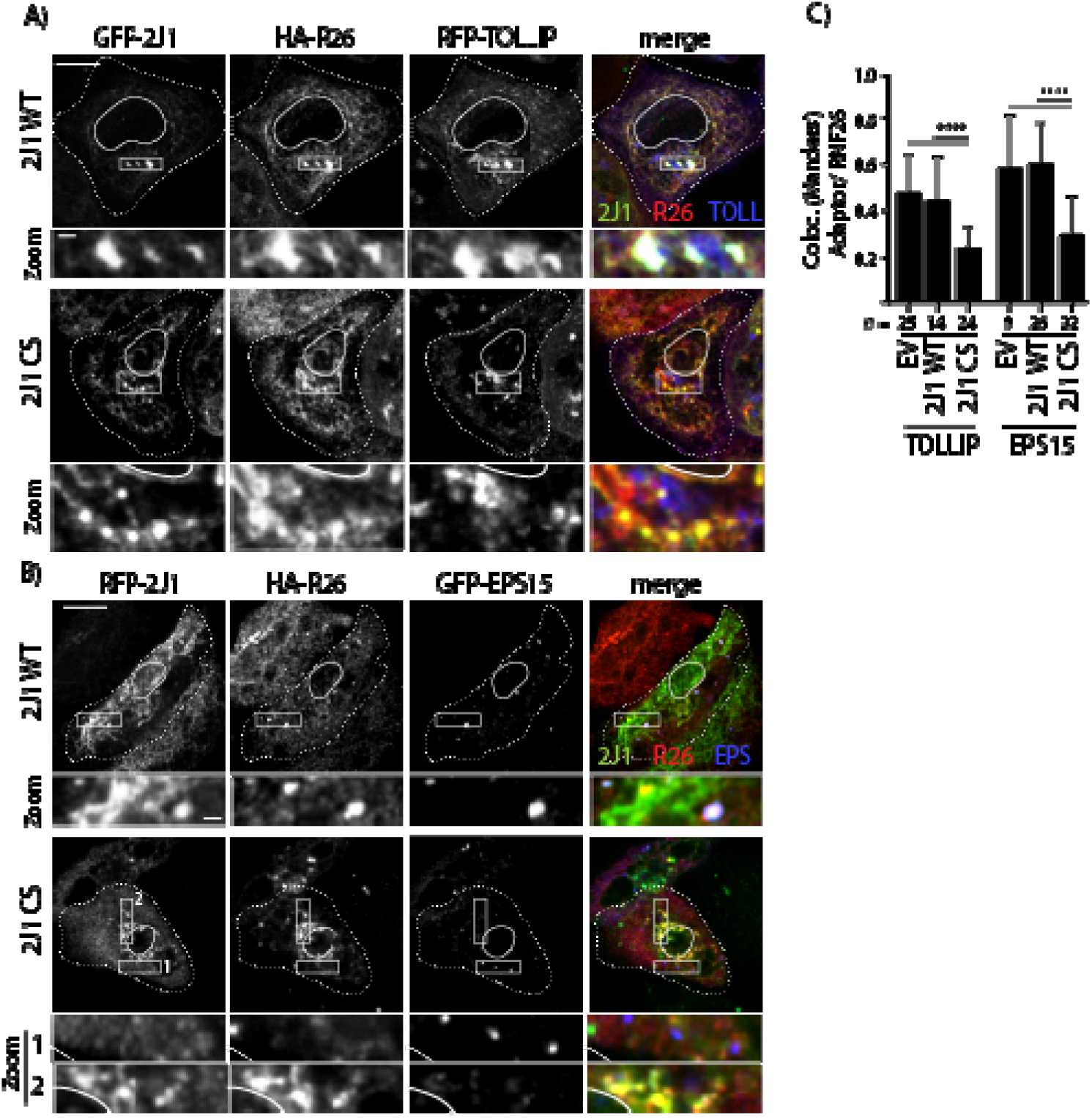
UBE2J1 catalytic activity is required for vesicle recruitment to RNF26. **A-B)** Tollip (A) or EPS15 (B) recruitment to RNF26 positive structures as a function of UBE2J1 activity. Confocal microscopy images of HeLa cells overexpressing RFP-TOLLIP (A), GFP-EPS15 (B), HA-RNF26 and WT or mutant (CS) GFP- (A) or REP- (B) UBE2J1. Shown are the separate channels and merged images of UBE2J1 (green), RNF26 (red) and TOLLIP/EPS15 (blue), and select zoom-ins. **C)** Quantification of overlap between HA-RNF26 and RFP-TOLLIP or GFP-EPS15 signal in HeLa cells overexpressing either UBE2J1 WT or mutant UBE2J1 CS, or empty vector (EV) control from (A) and (B). Scale bars = 10μm. Zoom scale bar = 1μm. Magnification identical for all images. ImageJ mean filter (1 pxl) was used to smoothen images. Cell and nuclear boundaries are demarcated using dashed and continuous lines, respectively. Number of cells analyzed per condition is given under each sample group. All statistical significance tested with Students’ T-test. * p<0.05; ** p<0.01; *** p<0.001; **** p<0.0001

### UBE2J1 promotes timely downregulation of EGFR

Spatiotemporal organization of the endosomal system translates into timely trafficking of activated EGFR to lysosomes for degradation (Jongsma et al., 2016). If acting upstream of RNF26, UBE2J1 is expected to also facilitate this process. To test this, we followed ligand-mediated trafficking and degradation of EGFR. Maturation of EGF-containing endosomes into the late compartments marked by CD63 was inhibited in HeLa cells depleted of UBE2J1 (Fig. 6A, B). As a consequence of impaired trafficking, these cells exhibited attenuated downregulation of stimulated EGFR, accompanied by a prolongation of the activated receptor state (Fig. 6C, D). Importantly, however, UBE2J1 silencing did not alter steady state levels of EGFR at the cell surface (Fig. 6E). These results underscore the physiological importance of UBE2J1 in the regulation of the endosomal system’s architecture and dynamics.

**Fig. 6:**
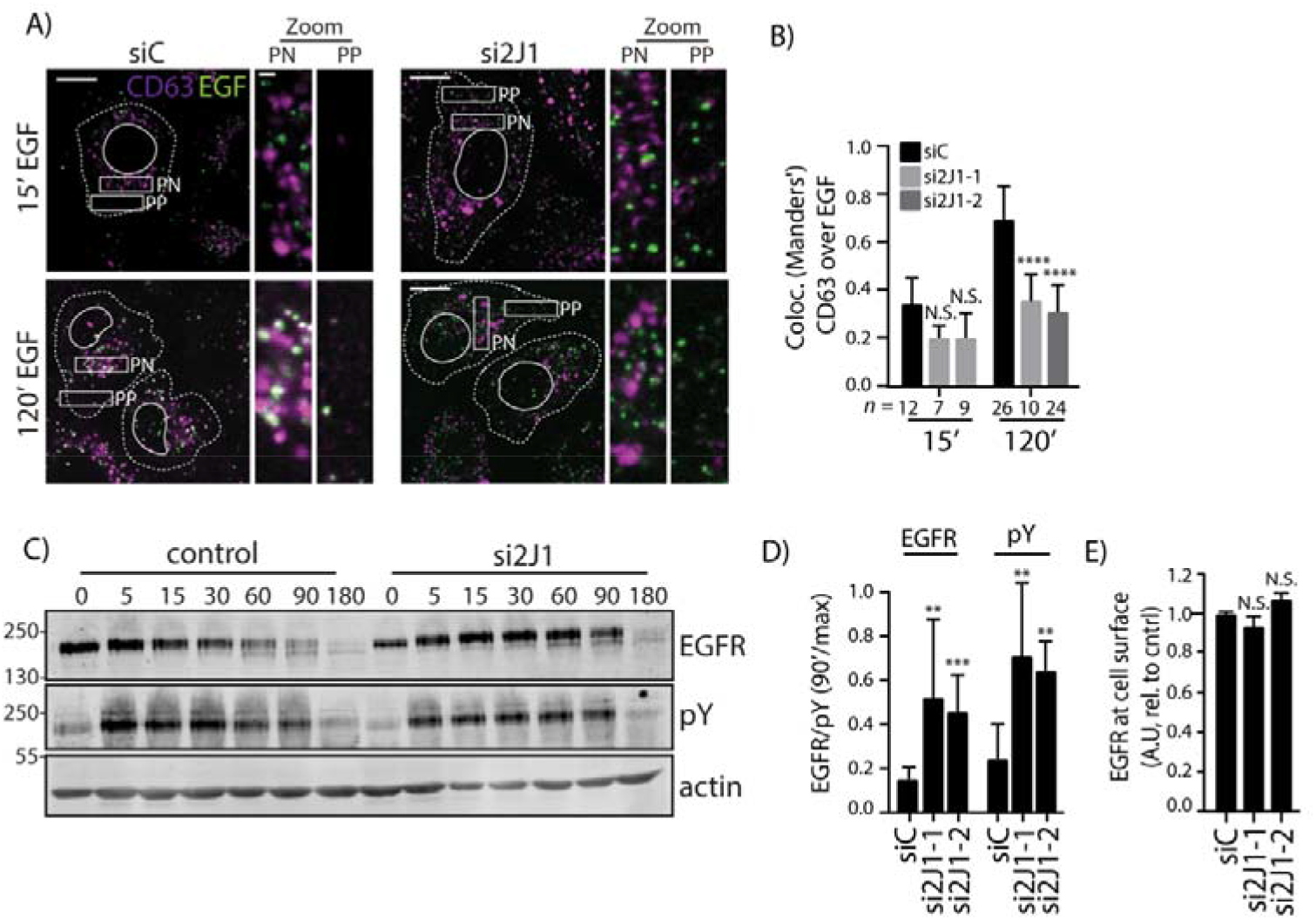
UBE2J1 is required for timely degradation and inactivation of EGFR. **A)** Trafficking of ligand-stimulated EGFR towards late endosomes in the presence or absence of UBE2J1. Representative confocal images show UBE2J1-1-depleted or control HeLa cells, stimulated with Alexa555-labeled EGF (100μg/mL) for 15 or 120 min before fixation and immunostaining against CD63. Shown are single focal plane images of EGF (green) and CD63 (magenta) at the indicated time points, and zooms of select perinuclear and peripheral areas. **B)** Colocalization (Manders’ overlap) of EGF-555 and CD63 in UBE2J1 silenced cells (si2J1-1 and si2J1-2) versus control cells at different time points, sampled from (A. Results show mean + SD of two independent experiments. **C)** Total and activated EGFR levels as a function of time in ligand-stimulated cells either or not depleted for UBE2J1-2. Shown are scans of Western blots stained for total and activated (pY) EGFR, as well as actin (loading control) levels at indicated time points following EGF stimulation in UBE2J1-2 silenced or control cells. **D)** Quantification of normalized amounts of total and activated (pY) EGFR at 90 min. after stimulation in (C). Quantification of total (EGFR, relative to t=0) and activated (pY, relative to full activation at t=15) EGFR levels at 90 minutes after EGF stimulation, normalized to actin levels. Shown is mean + SD of three independent experiments. **E)** EGFR expression as determined by flow cytometry (GeoMean) in cells transfected with siRNA or the control condition from (A). Shown is mean + SD of three independent experiments. Scale bar = 10μm. Zoom scale bar = 1μm. Magnification identical for all images. Cell and nuclear boundaries are demarcated using dashed and continuous lines, respectively. Number of cells analyzed per condition is given under each sample group. All statistical significance tested with Students’ T-test. ** p<0.01; *** p<0.001; **** p<0.0001

## Discussion

Endosomes rely on the ER to facilitate timely processing and selective delivery of cargoes for degradation. One aspect in which this manifests, is the architectural support the ER offers to the perinuclear endosome cloud—the cell’s hub for endosomal maturation and proteolysis (Neefjes et al., 2017). In this study, we implicate the ER-associated ubiquitin conjugating enzyme UBE2J1 in the perinuclear positioning of the endolysosomal system. Depletion of UBE2J1 disturbs the perinuclear vesicle cloud, where the bulk of these structures is normally retained in a low motility state. Consequently, motility patterns of endosomes and lysosomes throughout the cell are deregulated, and ligand-mediated trafficking of activated receptors to the proteolytic compartments is delayed. We find that UBE2J1 activity is a prerequisite for the recruitment of specialized endosomal adaptors to the ubiquitin-dependent positioning complex, assembled by the RING E3 ligase RNF26 at the ER membrane (Jongsma et al., 2016). We propose that UBE2J1 serves as an E2 for RNF26, and that endosomal phenotypes associated with UBE2J1 loss-of-function arise due to the inactivity of this perinuclear positioning pathway.

UBE2J1 has been extensively studied in the context of ER-associated degradation (ERAD) and stress recovery (Burr et al., 2011, Elangovan, Chong et al., 2017, Hagiwara et al., 2016, Lenk et al., 2002, Tiwari & Weissman, 2001), and its physiological roles in viral infection (Feng, Deng et al., 2018, Ma, Dang et al., 2015) and spermiogenesis (Koenig, Nicholls et al., 2014) are thought to be mediated through ERAD function(s). Our data reveal a new role for UBE2J1, which brings about the possibility that deleterious phenotypes associated with this enzyme’s dysfunction may also stem from defects in endosomal trafficking.

Several studies have shown that the nature of UBE2J1’s TMD determines its activity and stability (Claessen, Mueller et al., 2010, Yang M., 1997). Likewise, our data suggest that the interaction of UBE2J1 with RNF26 is stabilized mainly through their respective TMDs. Conversely, localization of the E2/E3 complex to a perinuclear subdomain of the ER is specified by the RING domain of RNF26. The resulting ER-embedded ubiquitylation complex modifies SQSTM1, thereby enticing ubiquitin-binding vesicle adaptors to dock at the ER membrane (Fig. 7). Thus, UBE2J1—an E2 enzyme primarily known for conducting poly-ubiquitylation—is now also implicated in mono and/or short-chain modifications by supporting RNF26 (Jongsma et al., 2016, Qin et al., 2014). In this regard UBE2J1 echoes a similar characteristic to that of its homologue UBE2J2/UBC6p (Weber, Cohen et al., 2016). Interestingly, we find that UBE2J2, also residing in the ER membrane (Wang, Herr et al., 2009, Weber et al., 2016), can similarly associate with RNF26, but does not support ubiquitin transfer to SQSTM1. In fact, overexpression of UBE2J2 appears to act as a dominant-negative for the UBE2J1/RNF26 pair. This pseudo-compatibility of UBE2J2 might relate to its cytoplasmic fragment, which is shorter than that of UBE2J1, perhaps making it more difficult for its UBC domain to reach substrates bound by the RING domain of RNF26. Single spanning TMD proteins, such as the aforementioned E2s, may be required for organization of large membrane-embedded protein complexes, as illustrated by interactions within the mitochondrial respiratory chain complex (Zickermann, Angerer et al., 2010). A similar organizational principle could hold for complexes at and/or within the ER membrane.

**Fig. 7:**
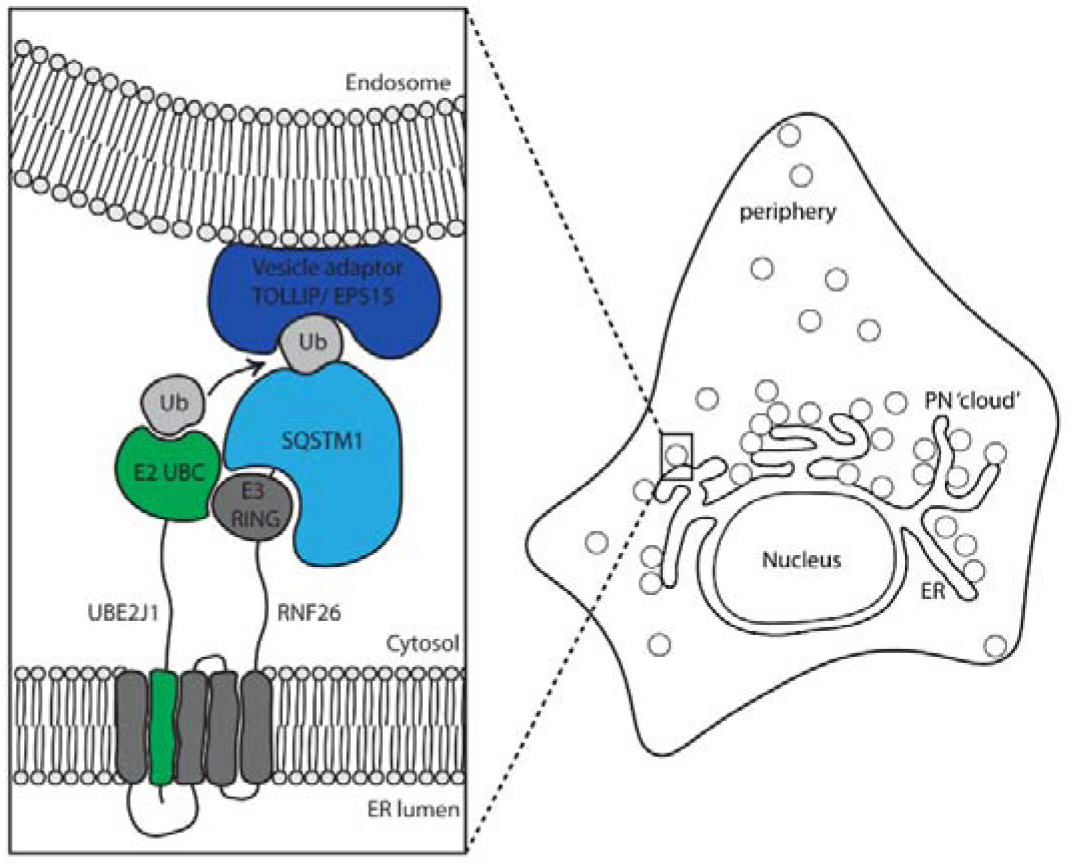
Model of the RNF26/UBE2J1 positioning complex in the ER membrane. The TMDs of RNF26 (dark grey) bind the single TMD of UBE2J1 (green) within the ER membrane to mediate close proximity of the RNF26 RING domain to the ubiquitin (light grey)-loaded UBE2J1 UBC domain that are both extending from the ER membrane into the cytosol. The activated UBE2J1/RNF26 complex ubiquitinates SQSTM1 (light blue) through mono-/short-chain linkage(s), which in turn serves as a platform for the binding of vesicle adaptors (dark blue) such as EPS15 or TOLLIP via their ubiquitin-binding domains. Hereby, vesicles bound by vesicle adaptors are recruited to the perinuclear endosomal cloud until release.

In recent years, numerous proteins have been identified to participate in the formation and regulation of ER-endosome contact sites, each regulating specific stages or transitions of endosomal trafficking. A maturing endosome can likely engage in multiple tethering interactions, and the composition and duration of its interactions with the ER could be influenced by maturation. Membrane proximity invoked by dynamic ubiquitin-mediated interactions, as described here, may enhance the strength/duration of ER-endosome membrane contact sites, providing a fundament for robust yet agile regulatory interactions. Finally, the UBE2J1/RNF26 endosomal positioning complex locates to defined perinuclear sites in the ER membrane that may be co-occupied by members of the ERAD family (Leitman et al., 2013). The close proximity of ERAD to lysosomes could then allow for swift alternate degradation of ERAD-resistant substrates.

## Materials & Methods

### Antibodies and reagents

*(Confocal microscopy)* mouse anti-EEA1 (1:200, mAb 610457, BD transduction laboratories), mouse anti-CD63 NKI-C3 (1:500, Vennegoor and Rumke, 1986), mouse anti-M6PR (1:100, ab2733, Abcam), mouse anti-TfR (1:100, Invitrogen 905963A), mouse anti-Ubiquitin (1:25, P4D1, sc-8017, Santa Cruz), rabbit anti-LAMP1 (1:200, Sino Biolocial), rabbit anti-VAP-A (1:40, 15272-1-AP, Proteintech), rat anti-HA (1:200, 3F10, Roche) followed by anti-Rabbit/Mouse/Rat Alexa-dye coupled antibodies (1:400, Invitrogen). LysoTracker FarRed was used to visualize lysosomes in live microscopy (1:2000, ThermoFisher). Alexa-568-coupled EGF (100ng/mL, Invitrogen) was used in endocytosis assays.

*(Western blotting)* mouse anti-HA (1:1000, HA.11, 16B12), rabbit anti-GFP (1:1000, (Rocha, Kuijl et al., 2009a), rabbit anti-RFP (1:1000, Rocha et al., 2019), mouse anti-RFP (1:1000, 6G6, Chromotek), rabbit anti-EGFR (1:1000, 06-847, Millipore), mouse anti-actin (1:20.000, AC15, Sigma), mouse anti-phosphotyrosine (1:1000, pY, 4G10, Millipore) followed by secondary Rabbit anti-mouse-HRP, sheep anti-rabbit-HRP (Invitrogen), or goat anti-rabbit or goat anti-mouse IRdye 680 (1:20.000) and IRdye 800 (1:10.000) antibodies (LiCor).

### Cell lines and culturing

MelJuSo cells (human melanoma), kindly provided by Prof. G. Riethmuller (LMU, Munich), were cultured in Iscove’s modified Dulbecco’s medium (IMDM) medium (Gibco). HeLa cells (CCL-2), and HEK293T cells were sequence verified cultured in DMEM medium (Gibco). All media were supplemented with 8% fetal calf serum (FCS, Sigma). All cell lines were cultured at 37□C and 5% CO2 and routinely tested (negatively) for mycoplasma.

### Constructs

RNF26 (and mutants), SQSTM1 and TOLLIP constructs, all expressed from C1/N1 vector series (Clontech), as well as HA-Ubiquitin and GFP-EP15 constructs (kind gifts from I. Dikic, Institute for Biochemie II, Frankfurt and O. Bakke, Dept. Bioscience, University of Oslo, respectively) have been previously described (Jongsma et al., 2016). UBE2J1 was subcloned between XhoI and BamHI sites of the C1-RFP and C1-GFP vector (Clontech). UBE2J1 truncations and mutants were created by standard (mutagenesis) PCR methods. UBE2J1 (MSD2TMD) and UBE2J1 (J2TMD) were created by fusing the cytoplasmic tail (aa1-282) of UBE2J1 to the TMD of MOSPD2 or UBE2J2 using NEBuilder HiFi DNA Assembly (NEB). pDEST17-UBE2J2 was a gift from Wade Harper (Addgene plasmid #15794) and UBE2J2-C93S was kindly provided by E. Wiertz (UMC Utrecht). From these plasmids, UBE2J2 and UBE2J2-C93S were subcloned into C1-RFP and C1-GFP using EcoRI and BamHI sites.

### siRNA transfections

For the initial E2 screen, pooled siRNAs were bought from Dharmacon (siGenome (M) series). Sequences of the siRNA oligos targeting RNF26 and UBE2J1, bought from Dharmacon, are shown in Table 1. For RNF26 silencing, we used siRNF26-1 unless stated otherwise. For UBE2J1 silencing, we used a pool of all three siRNAs unless stated otherwise. Gene silencing was performed in a 48 or 24 well plate (IF) or 12 well plate (WB) - reagent volumes were scaled up accordingly. In a 48 well plate, 25-32.5μL siRNA (for sequences, see table 1) was mixed with 25uL 1x siRNA buffer (GE Healthcare) containing 0.5uL Dharmafect 1 transfection reagent. The mix was incubated on a shaker at RT for 40 minutes before the addition of 7.000 HeLa or MelJuSo cells (and coverslips). Cells were cultured for three days before analysis. Non-targeting siRNA or reagent only was used as a negative control. UBE2J2 was silenced using siRNAs from the siGenome SMARTpool library (Dharmacon).

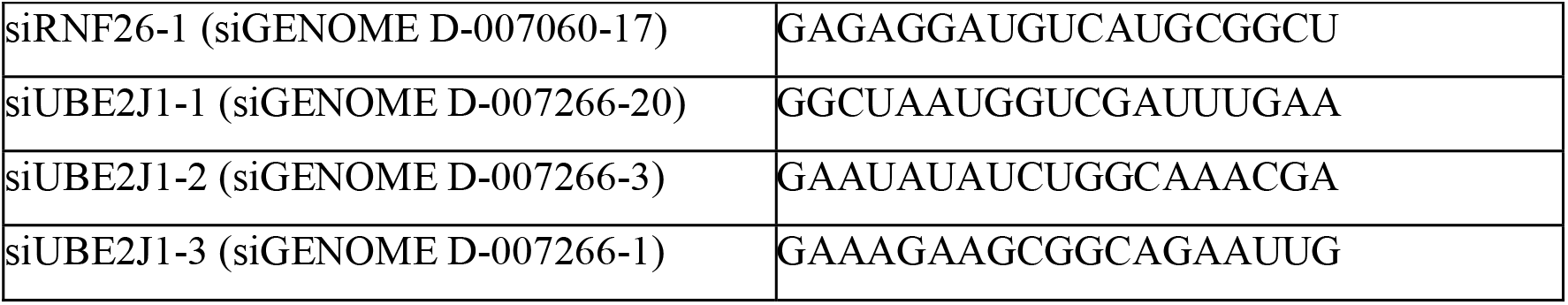

### DNA transfections

Cells were seeded in culture plates to reach approx. 70% confluency on the day of transfection. HeLa were transfected with Effectene (Qiagen) (200ng DNA per well of 24 well plate), according to the manufacturer’s protocol. MelJuSo cells were transfected using Extremegene HP (Roche) (500ng DNA per well in 24 well plate), according to manufacturer’s protocol. Cells were cultured overnight before analysis. HEK293 cells were transfected with PEI at a ratio of 3μg PEI per μg DNA in 200μL DMEM medium. After 15-30 min, the mix was added dropwise to the cells which were then cultured overnight before analysis.

### CRISPR/Cas9-mediated knockout

gRNA sequences targeting the UBE2J1 gene (GGGTCTCCATGGTGGGTCGC) were cloned into the BbsI site of PX440 (containing the Cas9 gene and a puromycin resistance gene). This plasmid was transfected into HeLa cells and the next day, cells were selected with 200ug/mL puromycin for 3 days. Then, cells were diluted and cultured in a 15cm dish, allowing well separates colonies to grow. These were isolated, expanded, and analyzed for loss of UBE2J1 by Western blot.

### EGFR degradation

Ligand-mediated turnover was assayed as previously described (Berlin, Higginbotham et al., 2010) using 100 ng/mL EGF. Receptor levels were quantified at each time point relative to actin levels and expressed as a fraction of EGFR at t = 0 min. Receptor activation and downstream signaling was expressed relative to the maximum activation (t = 15 min).

### Immunofluorescence confocal microscopy

Cells grown on coverslips (Menzel Gläser) were fixed with 3.7% paraformaldehyde, washed three times with PBS, permeabilized with 0.1%TX100 for 10 min and blocked in 0.5% BSA for one hour. Next, coverslips were incubated with primary antibodies in 0.5% BSA for 1hr at RT, washed and incubated with Alexa-labeled anti mouse/rabbit/rat secondary antibody or streptavidin. After washing, coverslips were mounted on glass slides with ProLong Gold with DAPI (Life Technologies). Samples were imaged with a Leica SP8 confocal microscope equipped with appropriate solid-state lasers, HCX PL 63 times magnification oil emersion objectives and HyD detectors (Leica Microsystems, Wetzlar, Germany). Data was collected using 2048 x 2048 scanning format with line avering without digital zoom, or 1024 x 1024 scanning format with digital zoom in the range of 1.0-2.0 with line averaging. Quantification of endosome positioning was performed as previously described with minor alterations (Jongsma et al., 2016; Sapmaz et al., 2019). In short, fluorescence intensities (above automated background threshold) were measured along a straight line ROI (regions of interest) drawn from the border of a cell’s nucleus (fractional distance = 0) to the plasma membrane (fractional distance = 1.0) using the line profile tool in the LAS-AF software, and their absolute distance to the border of the nucleus was expressed relative to the total length of the line. Fractional distances are reported in scatter plots along with the median distance value (red line) within the sample and the total number of cells analysed.

### Live microscopy

For live microscopy, cells were seeded in 4-chamber live cell dishes and imaged under conditions of 37°C and 5% CO2 with a Leica SP8 WLL confocal microscope. Data was collected using 63x oil immersion objectives and 1.5x magnification in a 1024 x 1024 scanning format at 0.58 frames/sec with line averaging. Tracking of lysotracker-positive vesicles was performed using TrackMate for FiJi. FiJi was also used for post-collection image processing.

### Co-immunoprecipitations

HEK293T cells were lysed in 1% DMNG buffer (150mM NaCl, 50mM Tris-HCl pH 7.5, 5mM MgCl2, 1% DMNG, protease inhibitors (Roche diagnostics, EDTA free) for 90 min, rotating at 4°C. After 15 min 20,000x *g* centrifugation, post-nuclear lysates were incubated with GFP-TRAP beads (Chromotek) and rotated for 90 min at 4□C before subsequent immunoisolation with RFP-TRAP beads to acquire input samples. Beads were washed four times in 0.2% DMNG lysis buffer. Samples were boiled for 10 min in 2x Laemmli buffer prior to analyses by SDS-PAGE and Western blotting.

### Ubiquitination assays

HEK293 T cells were lysed in 0.5% TX-100 lysis buffer (150mM NaCl, 50mM Tris-HCl pH 7.5, 5mM MgCl2, 0.5% (v/v) TX-100, 20mM NEM, protease inhibitors (Roche diagnostics, EDTA free)). Nuclei were lysed by adding 1:4 SDS buffer (2% SDS, EDTA) and samples were sonicated (5×1s pulses, 80% power, Fisher Scientific). Samples were diluted to 0.2% SDS with TX-100 lysis buffer and centrifuged for 20 min at 20,000 x *g.* After spinning, samples were incubated with GFP-TRAP beads (Chromotek) for 3hrs at 4°C. Beads were washed 3 times with 8M Urea and 1% SDS in PBS, and 1 time with 1% SDS in PBS before elution in 2x Laemmli sample buffer by boiling for 10 min.

### SDS-PAGE and Western blotting

Samples were separated by 8% (ubiquitination assays) or 10/12% (regular lysates, CoIPs) SDS-PAGE and transferred to nitrocellulose or PVDF membranes in ethanol-containing transfer buffer at 300mA for 2-3 h. The membranes were blocked with 5% milk/PBS before incubation with primary antibody diluted in blocking buffer for 1hr at RT. After washing twice with PBS/0.1% Tween-20, proteins were detected with secondary antibodies. Depending on the secondary antibody, detections were performed by incubation with ECL reagent (SuperSignal West Dura Extended Duration, GE Healthcare) or directly imaged with an Odyssey Fx laser scanning fluorescence imager.

### Flow cytometry

For detection of cell surface EGFR, cells were trypsinized and suspended in FACS buffer (2% FCS in PBS) and stained with PE-conjugated anti-EGFR for 30 min on ice. After two washes with ice-cold FACS buffer, cells were fixed in FACS buffer containing 0.1% PFA until analysis by a FACS Calibur flow cytometer (BD Biosciences).

## Acknowledgements

This work was supported by an ERC Advanced Grant ERCOPE to JN.

## Author Contributions

Conceptualization, T.C. and I.B.; Methodology, T.C. and I.B.; Formal Analysis: T.C.; Investigation, T.C. and M.J.; Writing – Original Draft, TC and J.N.; Writing – Review & Editing, I.B. and J.N.; Visualization: T.C.; Funding Acquisition, I.B. and J.N.; Supervision, I.B. and J.N.

## Supplemental movie 1

Time lapse of LysoTracker-positive vesicle movement in control vs UBE2J1 depleted MelJuSo cells. Cells were transfected with siRNAs three days before imaging at 1.62 seconds per frame for 105 frames.

---

**Fig. S1:**
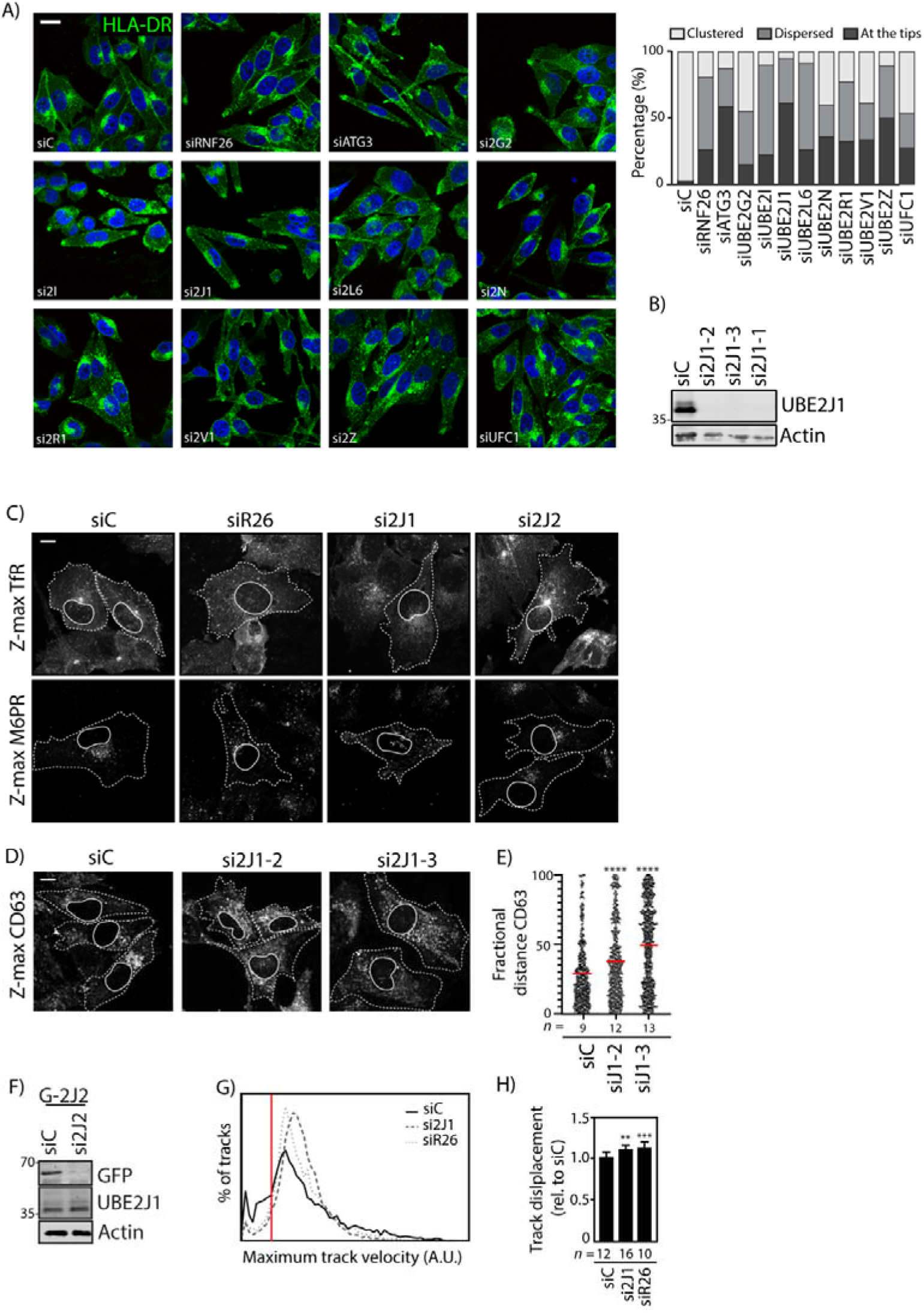
Screen for E2 enzymes affecting the architecture of the endolysosomal system and validation of UBE2J1 loss of function phenotype. Related to Fig. 1. **A)** Distribution of late endosomal compartments in response to siRNA-mediated depletion of the indicated E2 enzymes or RNF26 (positive control). Representative confocal images of MelJuSo cells transfected as indicated and stained against late endosomal MHC class II cargo (HLA-DR, green) and nuclear DAPI (blue). Bar graph reports vesicle distribution quantified by categorizing cells as harboring a clustered (white), dispersed (light gray), or “tip” (dark gray) phenotype, 20-50 cells analyzed per condition. Scale bar = 20μm; same magnification for all subfigures. **B)** Western blot analysis of UBE2J1 knockdown efficiency in MelJuSo cells transfected with three different siRNAs. Endogenous UBE2J1 levels are shown with actin as loading control. 35kD molecular weight marker indicated. **C)** Distribution of recycling endosomes (TfR^+^), TGN vesicles (M6PR^+^) and MelJuSo cells depleted of RNF26, UBE2J1 and UBE2J2 as assayed by confocal microscopy. Maximum Z- proj ections are shown with cell boundaries depicted in dashed lines and nucleus (deduced from DAPI stain) with continuous line. **D)** Distribution of late endosomes (CD63^+^) in MelJuSo cells in response to depletion of UBE2J1 with two other siRNAs. Representative maximal projection confocal images of MelJuSo cells transfected as indicated and stained against CD63. **E)** Vesicle localization expressed as fractional distance of fluorescent pixels along a straight line drawn through the endosomal cloud from the nuclear edge (0) to the plasma membrane (1.0) analyzed from samples in (S1D). Red line: mean, n=2 independent experiments. **F)** Western blot analysis of UBE2J2 knockdown efficiency in HeLa cells overexpressing GFP-UBE2J2. Shown are overexpressed GFP-UBE2J2 levels and endogenous UBE2J1 levels with actin levels as loading control after transfection with pooled siRNAs targeting UBE2J2. Position marker proteins indicated. **G)** Total track velocity distribution expressed as histogram of pooled samples from (1E) in control cells versus those depleted of either UBE2J1 or RNF26. Red line: cutoff to define “low velocity tracks” in (1D). **H)** Quantification of vesicle displacement in control MelJuSo cells (siC) or those depleted of either UBE2J1 (si2J1-1) or RNF26 (siR26). n=2 independent experiments. Shown is mean+SD. Scale bars = 10μm. Magnification identical for all images. Cell and nuclear boundaries are demarcated using dashed and continuous lines, respectively. Number of cells analyzed per condition is given above each bar/scatter. All statistical significance tested with Students’ T-test. **** p<0.0001.

**Fig. S2:**
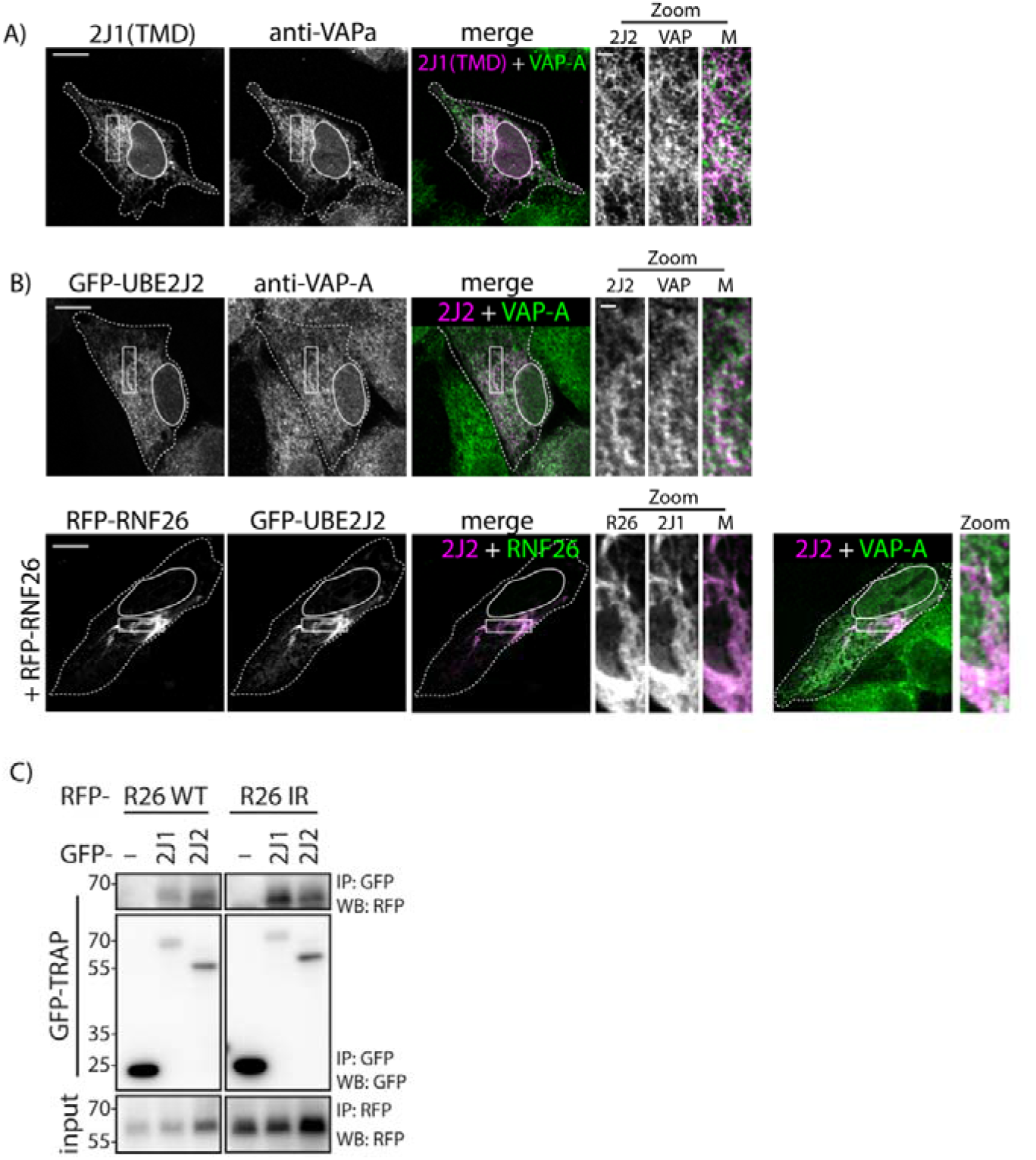
Localization of UBE2J1 TMD in the ER membrane and interaction and relocalization of UBE2J2 to a perinuclear ER region. Related to fig. 2 and 3. **A)** Localization of the GFP-UBE2J1 TMD segment in the ER. Representative confocal image of MelJuSo cell expressing GFP-UBE2J1, with the ER labelled by VAP-A. Rectangular zoom-ins highlight select regions of indicated channels. **B)** Localization of GFP-UBE2J2 in MelJuSo cells with or without co-expression of RFP-RNF26. VAP-A staining indicate position of ER. Overlays of either UBE2J2 and VAP-A or RNF26 are shown. Rectangular zoom-ins highlight select regions of indicated channels. **C)** Interactions between RFP-RNF26 or its inactive I382R (IR) mutant and GFP-UBE2J1 or - UBE2J2, as assayed by co-IP. For input samples, post-IP lysate was subjected to a second IP with an excess of RFP-TRAP beads. Position of the marker proteins indicated. Representative gel out of three independent experiments. Scale bars = 10μm. Zoom scale bars = 1μm. Cell and nuclear boundaries are demarcated using dashed and continuous lines, respectively. Magnification identical for all images.

**Fig. S3:**
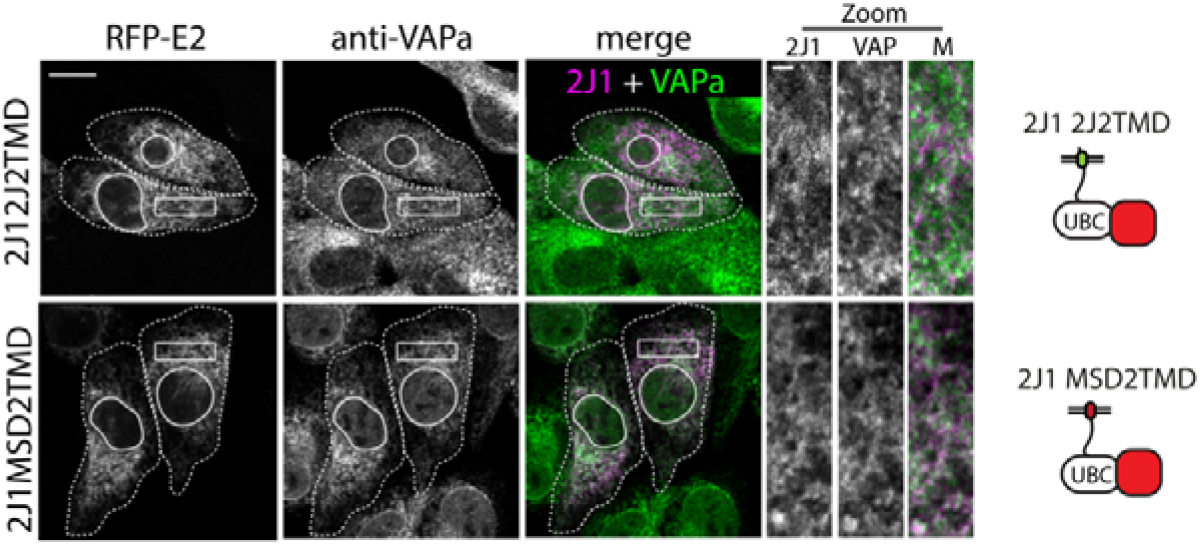
Correct localization of the UBE2J1 TMD mutants in the ER. Related to Fig. 4. Localization of RFP-UBE2J1 mutants 2J2TMD and MSD2TMD (from Fig. 4G) in the ER. Cells were immunostained from the ER marker VAP-A. Single focal plane images are shown. Scale bar = 10μm. Zoom scale bar = 1μm. Magnification identical for all images.

